# Optimal decoding of NFkB signaling dynamic

**DOI:** 10.1101/595272

**Authors:** Alok Maity, Roy Wollman

**Affiliations:** Institute for Quantitative and Computational Biosciences, University of California, Los Angeles.; Departments of Integrative Biology and Physiology and Chemistry and Biochemistry, University of California UCLA.

## Abstract

The encoder/decoder paradigm suggests that signaling networks transform information about the extracellular environment into specific signaling patterns that are then read by downstream effectors to control cellular behavior. Previous work used information theoretical tools to analyze the fidelity of encoding using dynamic signaling patterns. However, as the overall fidelity depends on both encoding and decoding, it is important to consider information loss during signal decoding. Here we used NFkB signaling as a model to understand the accuracy of signal decoding. Using a detailed mathematical model we simulated realistic NFkB signaling patterns with different degrees of variability. The NFkB patterns were used as an input to a simple gene expression model. Analysis of information transmission between ligand and NFkB and ligand and gene expression allow us to determine information loss in both encoding and decoding steps. Information loss could occur due to biochemical noise or due to lack of specificity in decoding response. We found that noise free decoding has very little information loss suggesting that decoding through gene expression can preserve specificity in NFkB patterns. As expected, information transmission through a noisy decoder suffers from information loss. Interestingly, this effect can be mitigated by a specific choice of decoding parameters that can substantially reduce information loss due to biochemical noise during signal decoding. Overall our results show that optimal decoding of dynamic patterns can preserve ligand specificity to maximize the accuracy of cellular response to environmental cues.

**Synopsis:** The fidelity of signal transduction depends on the accurate encoding of ligand information in specific signaling patterns and the reliable decoding of these patterns by downstream gene expression machinery. We present an analysis of the accuracy of decoding processes in the case of the transcription factor NFkB. We show that noiseless decoding can preserve ligand identity with minimal information loss. Noisy decoding does result in information loss, an effect that can be largely mitigated by choice of optimal decoding parameter values.

- Decoding of dynamic signaling patterns by a simple gene model can preserve most of the information about ligand identity.
- Noisy decoding will result in information loss, but this effect can be mitigated by the optimal choice of decoding parameters.
- Improvement in decoding is a result of decreased variability in gene expression patterns.

## Introduction

The ability of cells to respond to environmental changes is key to their function. A ligand binding to a receptor initiates a cascade of biochemical transformations of cellular kinases, phosphatase, and other enzymes that connect the receptor to downstream effector, often to change gene expression patterns (Lim *et al*, 2014). Historically, these cascades of events were divided into distinct pathways. The pathway paradigm was appealing because it was easy to understand how information about ligand identity and abundance is preserved. However, as more and more signaling interactions were discovered, it became clear that the linear signaling pathway paradigm is insufficient in describing the complexity of how information propagates between receptors and effectors. The complexity of signaling network with a large degree of crosstalk and feedback shifted the paradigm from pathway to network-centric view (Bhalla & Iyengar, 1999). However, unlike an isolated signaling cascade, the network view poses a challenge, how does the information about ligand identify and concentration is preserved in the interaction network?

A plausible solution for the specificity challenge in signaling networks is based on the concept of signal encoding and decoding (Behar & Hoffmann, 2010; Purvis & Lahav, 2013). Information about extracellular events undergoes multiple transformations. Initially, information on ligand identity and abundance is transformed, i.e. encoded, into a specific signaling activity pattern.

Subsequently, downstream effectors such as transcription factors transform, i.e. decode, the specific signaling activity patterns to a specific cellular response (Dolmetsch *et al*, 1998; Hoffmann *et al*, 2002; Hao & O’Shea, 2012; Batchelor *et al*, 2011). The encoding/decoding view is useful since it explains how information can be preserved despite the complex many to many relationships between receptors and signaling nodes. However, to what degree does biochemical noise and cellular variability limits reliable encoding and decoding is unclear.

Information theory can be used to assess the quality of signaling networks fundamental function, to reliably transmit specific information about ligand concentration to downstream effectors, allowing the cell to adjust its physiological state to changing conditions (Uda & Kuroda, 2016; Tkačik & Bialek, 2016; Levchenko & Nemenman, 2014; Bowsher & Swain, 2014; Waltermann & Klipp, 2011; Nandagopal *et al*, 2018; Wilson *et al*, 2017). Existing biochemical variability that occurs at multiple timescales can have an adverse effect on the quality of information transmission. Using information theoretical tools, one can probe the operational quality of a signaling network in a quantitative manner by measuring the information transmission capacity of a network through a series of input/output measurement. Mutual information analysis can quantify the degree of overlap between cellular responses to multiple distinct inputs and thereby it is a good descriptor of the accuracy of information transmission through a signaling network. These tools have been applied to signaling networks showing that indeed information loss occurs due to biochemical noise during encoding step (Selimkhanov *et al*, 2014; Uda *et al*, 2013; Cheong *et al*, 2011; Hansen & O’Shea, 2015; Potter *et al*, 2017).

Within the encoding/decoding paradigm, information transmission is only as good as the weaker of the two. If decoding is noisy and inaccurate, a high-quality encoding is of little use to the cell. Therefore, the question of how accurate is the decoding component of the network is paramount to our understanding of the overall performance characteristics of signaling networks. Despite this importance, only a few studies have addressed this question. Work by Hansen et al (Hansen & O’Shea, 2015) analyzed the degree of gene expression accuracy with regards to oscillatory signal and amplitude. While useful, it is unknown if cells rely on amplitude and frequency as the key features. In a more physiological setting, Lane et al measured the correspondence in a single cell between NFkB dynamics and resulting gene expression (Lane *et al*, 2017). They were able to demonstrate that indeed the overall patterns of signaling dynamics are decoded into distinct expression patterns. However, technical limitation and small sample size preclude an analysis of the reliability of the decoding processes. Therefore, there is a gap in our understanding of the degree of accuracy of information decoding by cellular effectors.

Here we address this gap by analyzing the decoding quality of a simple gene expression model. We utilize NFkB signaling, as it is a system where it’s encoding and decoding dynamics have been analyzed in depth (Cheong *et al*, 2011; Selimkhanov *et al*, 2014; Uda *et al*, 2013; Habibi *et al*, 2017; Lee *et al*, 2014). We first generate simulated encoding data for multiple ligands that activate NFkB and ask how good is the decoding of these encoded dynamic profiles. We show that without decoding noise, decoding close to optimal can by a large range of parameter values. However, when decoding noise is considered, close to optimal decoding can still be achieved by specific optimal decoding parameters. Overall our finding demonstrates that information decoding by gene expression machinery can be achieved to allow cells to accurately respond to environmental changes.

## Results and Discussion

We first constructed mathematical models that represent the encoding and decoding steps in NFkB signaling (Fig 1). The first model (Fig 1A) captures the encoding step and the transformation of ligand identify into the dynamics of NFkB. The second model (Fig 1B) focuses on gene expression decoding of NFkB dynamics. As the goal of the encoding model was to capture the specific aspects of NFkB dynamics we opted to use a highly detailed model with 95 reactions of 48 reactants and 127 parameters. The model connects five different receptors: TNFR, TLR4, TLR3, TLR9, and TLR1/2 through the key downstream pathways of MyD88 and TRIF to the IKK modules that controls NFkB dynamics (Cheng *et al*, 2015). The model generates realistic simulated time series up to 8 hours that match experimental data for five different ligands (Taylor et al, 2019). Given the scope of the model and the fact that includes multiple inputs to the same core network we focused on the question of ligand specificity and tested one ligand concentration for each condition (i.e., TNF = 10ng/ml, LPS = 20ng/ml, PolyIC = 30μg/ml, CPG = 10μM, Pam3CSK = 100ng/ml). The ligand concentrations were chosen to be close to the corresponding receptor Kd. We focused on the question of ligand specificity The key benefit of using a mathematical model to capture NFkB dynamic profile is that it allows us to control the degree of encoding noise, a manipulation that is technically impossible to perform experimentally. Furthermore, the analysis avoids the need to deal with experimental and technical measurement errors. The second model (Fig 1B) captures the decoding step. Unlike the encoding model that aims at highly realistic details description of the underlying system, the decoding model is very simple with a single differential equation and six parameters. We chose a simple model to allow us to capture the essence of the decoding step without unnecessarily increasing the number of parameters. Using the two models in tandem we can simulate how cells respond to extracellular stimuli through both NFkB signaling dynamics and resulting gene expression at multiple degrees of biological extrinsic noise.

**Figure 1.**
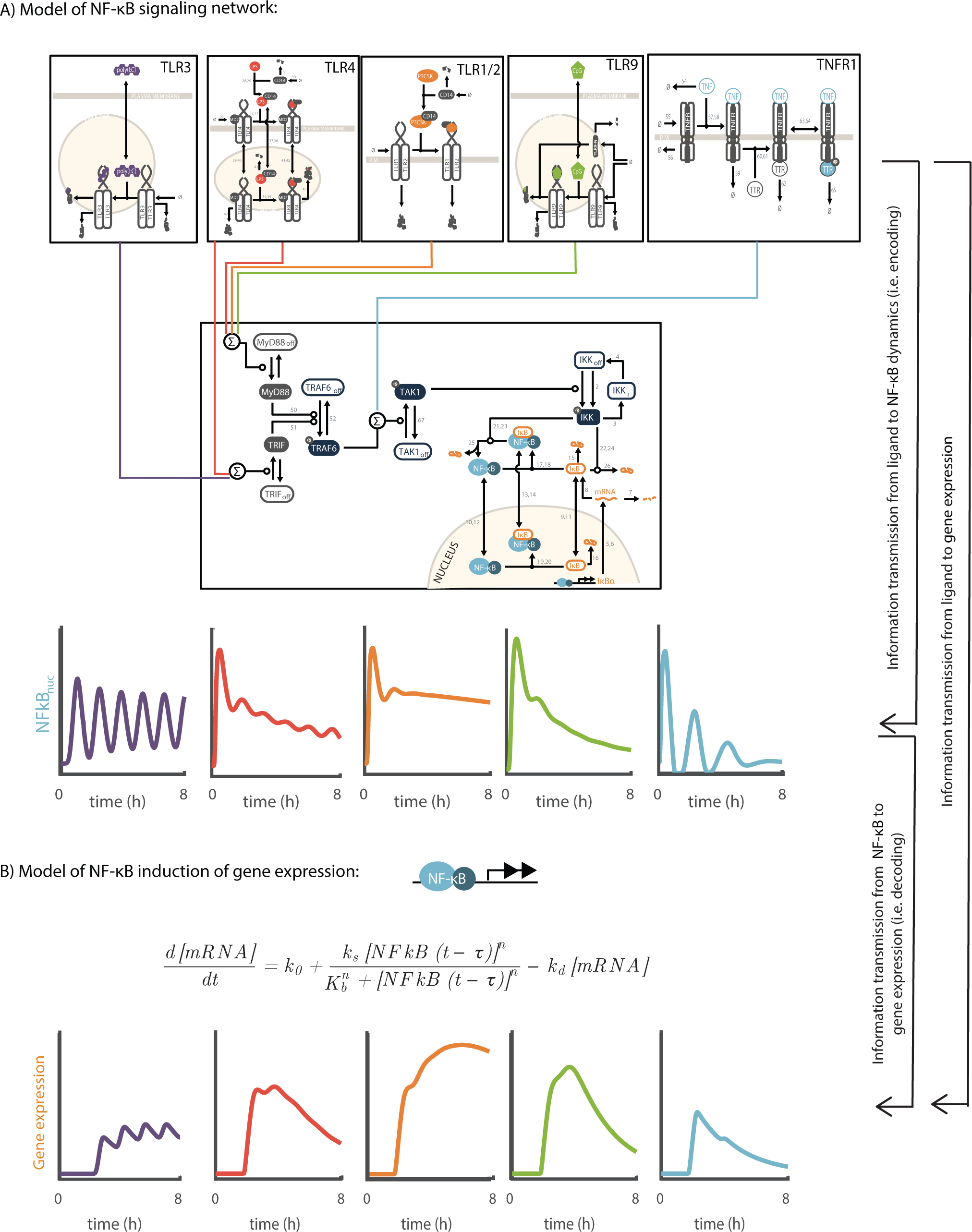
Models of encoding and decoding in the NfkB network. **A**. The encoding model of NFkB signaling network. The model includes five distinct receptor modules (TLR3, TLR4, TLR1/2, TLR8, TNFR1) that feed at three distinct points (MyD88, TRIF, and TAK1) into the core IKK/NFkB module. The dynamic interaction in network results from the mapping between each receptor module and a dynamic activation profiles of the transcription factor NFkB. **B**. The decoding model is highly simplified model represented by a single ordinary differential equation that uses the nuclear concentration of NFkB over time as an input and produces the gene expression pattern as an output. The decoding model effectively maps the dynamic of NFkB into dynamic gene expression profile.

To analyze the effect of cellular heterogeneity on the encoding and decoding steps we estimated the effect of biochemical variability on the accuracy of signal transduction using an information theoretic approach. The dominant source of variability in many signaling network (Toettcher *et al*, 2013; Yao *et al*, 2016; Selimkhanov *et al*, 2014) and in NFkB specifically (Hughey *et al*, 2015) is differences between cells in their underlying cell state (e.g. protein concentration, organelle structure, etc) between cells. To capture this effect we repeated the simulation of the encoding model 1000 times per condition with parameter drawn from a log-normal distribution centered around the reference value with different degrees of the coefficient of variation. We tested a range of variability magnitudes from 10-35% and at each condition estimated the mutual information between ligand identity and NfkB dynamics (Figure 2A, cyan line). As expected, the accuracy of encoding step depended on the degree of biochemical variability. At low values (10%) there was very little information loss compared to the maximum possible value of log_2_(5) ~ 2.3. As expected, as the magnitude of variability increased the signaling accuracy decreased.

**Figure 2.**
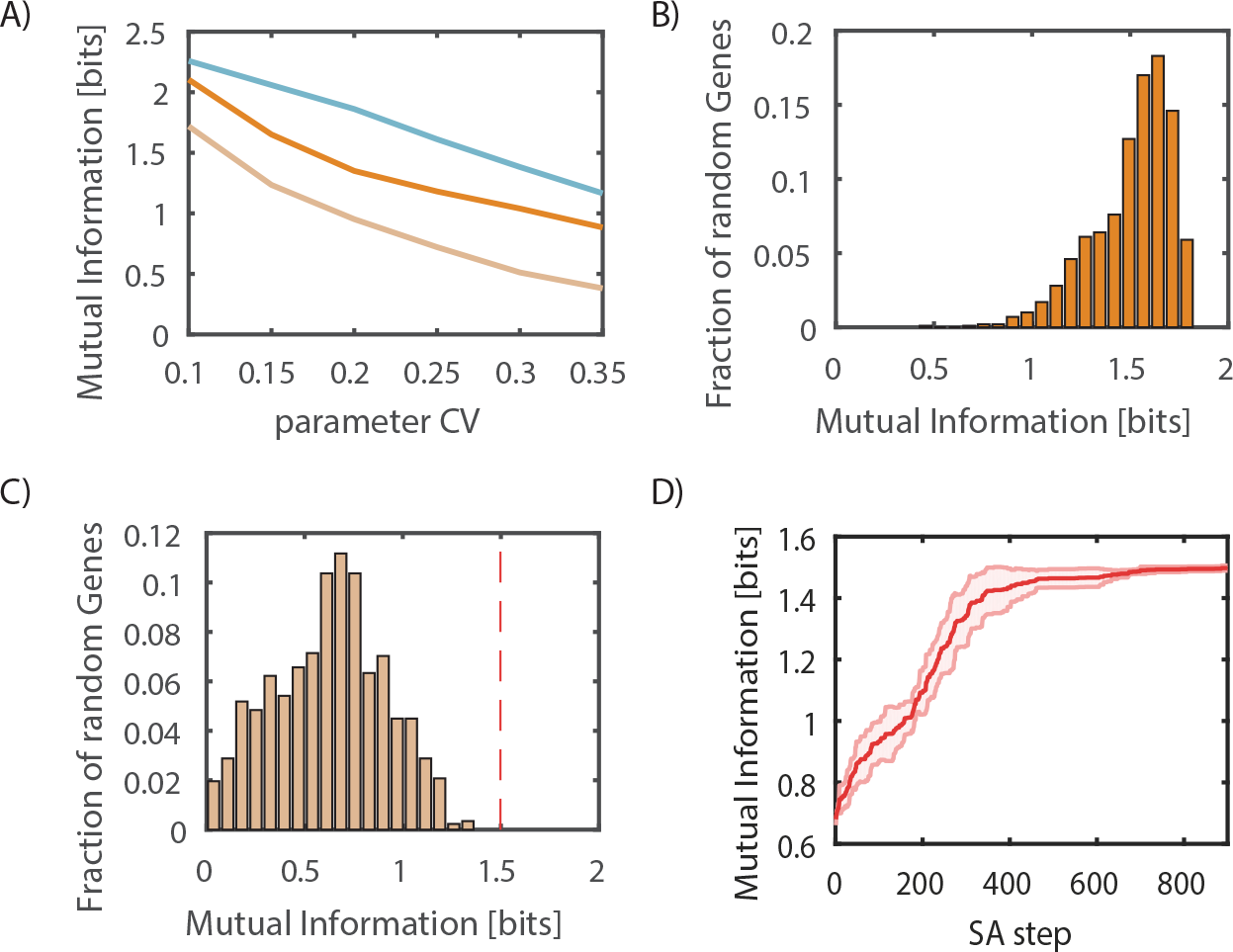
Information loss in NfkB dynamic encoding and decoding. **A.** Mutual information as a function of parameter variability that exists in the encoded in NFkB dynamics (cyan), and noiseless (dark orange) and noisy (light orange) decoding of NFkB by a simple gene expression model. **B** and **C** distribution of mutual information values in noiseless (**B**) and noisy (**C**) decoding of NFkB dynamics by a simple gene expression model. The distribution comes from the inference of mutual information after sampling different model parameters. In each case, 1000 parameter values are sampled from a log-normal distribution centered around reference values with a CV of 25%. The mutual information achieved after optimization of the parameter values of the decoding model is shown as a red dashed line. **D**. Optimization of parameter values using heuristic simulated annealing show convergence of mutual information values. The red line shows the average and shading the standard deviation from 4 repeated optimization runs.

Next, we used the simulated trajectories as input to the decoding model and asked how much additional information loss will occur during decoding. To separate the effect of biochemical variability during encoding and decoding we first tested decoding without the addition of biochemical variability in decoding steps themselves. To make the analysis independent of the specific parameter values of encoding we sampled 1000 different parameter values. We found that noise free decoder is able to capture most of the information that within NfkB dynamics (Fig 2AB). This indicates that the five tested ligands generate sufficiently distinct NfkB activation profiles such that a simple model of gene expression is sufficient to generate distinct gene expression profiles that provide sufficient information capture the ligand identity to a similar degree that it exists in NFkB dynamics.

The analysis above takes into account noise within the encoding step but ignores the existence of noise within the decoding step. As it is likely that biochemical variability plays a role in the decoding of signal we tested the overall information transmission fidelity in the presence of noise in the decoding step. Addition of biochemical variability into the decoding step resulted in additional loss of information. This was true for large ranges of parameter values for the decoding model (Fig 2C).

To maximize signaling fidelity, information loss should be minimized. We tested whether the information loss we observe during the decoding step is an unavoidable property of the system, or whether it is possible to achieve higher fidelity in decoding with optimized parameters. We performed heuristic optimization of decoding parameters with the objective function of maximizing mutual information between ligands and gene expression dynamics (Figure 2D). Interestingly, with optimal parameters, there was little loss of information during the decoding step (Fig 2C red dashed line) despite the existence of decoding noise. Repeating the optimization multiple time identified similar parameter values. Overall these results indicate that careful choice of the population average decoding parameters can mitigate the effect of decoding variability around these reference values. These results suggest that cell can potentially adopt an optimal decoding strategy to maximize the information extraction from the signal encoded by NFkB dynamics.

We next wanted to better understand the identified optimal strategy by answering two questions related to the optimal solutions: 1. What changes in gene expression patterns cause it to increase its information about ligand identity and 2. What model parameters are changed and do these changes contribute to the increase in decoding accuracy?

To assess what changed about the system after optimization we examined the dynamic trajectories of NFkB and resulting gene expression patterns (Figure 3). A strength of mutual information analysis is that it incorporates both differences in the overall separation between average response to ligand and the variability within a response to a single ligand into the same framework. Analysis of the gene expression pattern before and after optimization indicates that the major change in gene response after optimization is a reduction in response variability per each ligand (Figure 4D) and not from an increased separation of the typical response to all ligands (Figure 4E).

**Figure 3.**
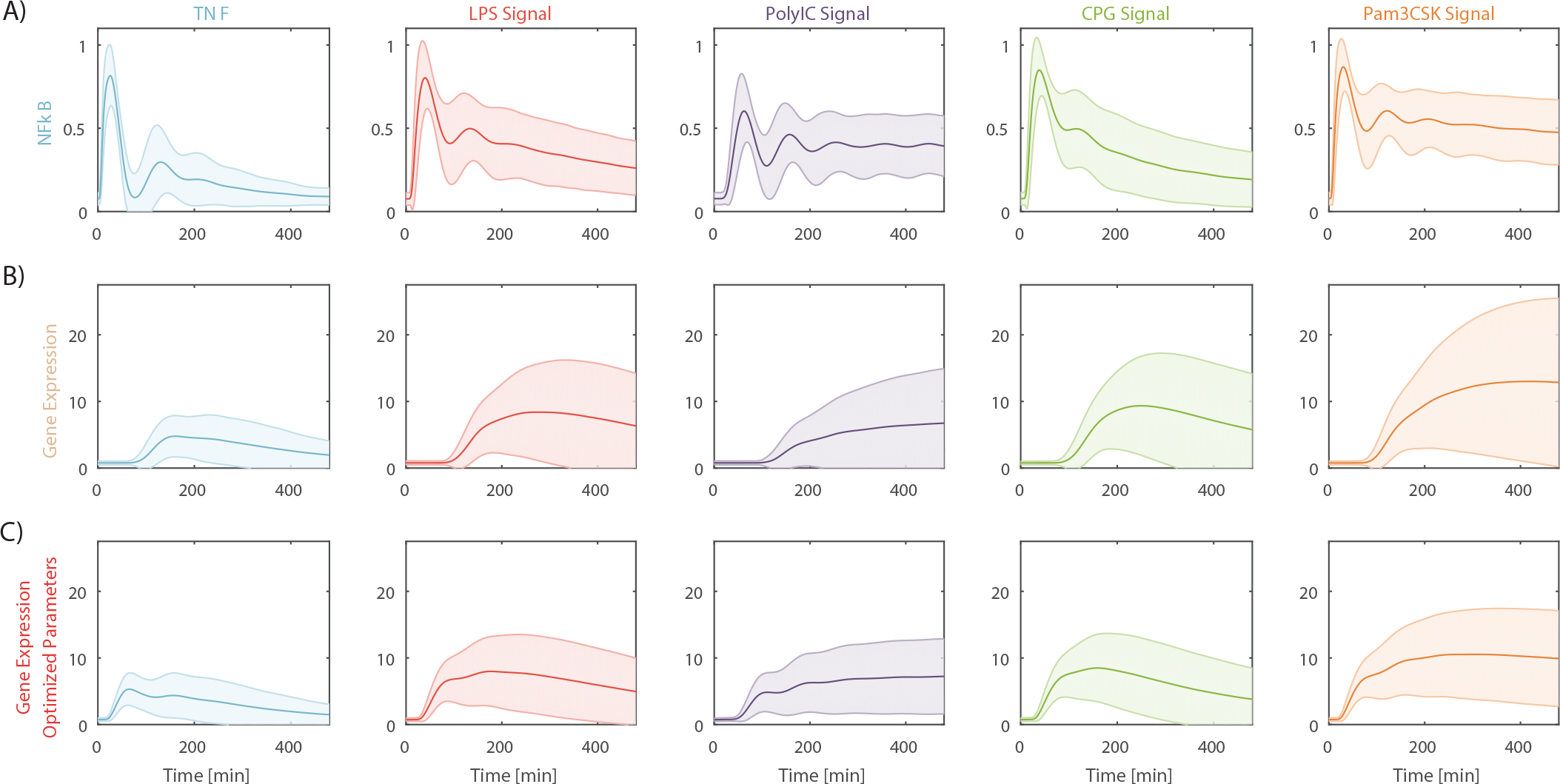
Simulation of NfkB dynamics and resulting gene expression. Average (center line) and standard deviation (shading) of NFkB dynamic (**A**), noisy gene expression decoder (**B**) and noisy gene expression decoder centered around optimal parameter values (**C**). Each condition was simulated 1000 times with parameter of both encoding and decoding models sampled from a log-normal distribution with a CV of 25%.

**Figure 4.**
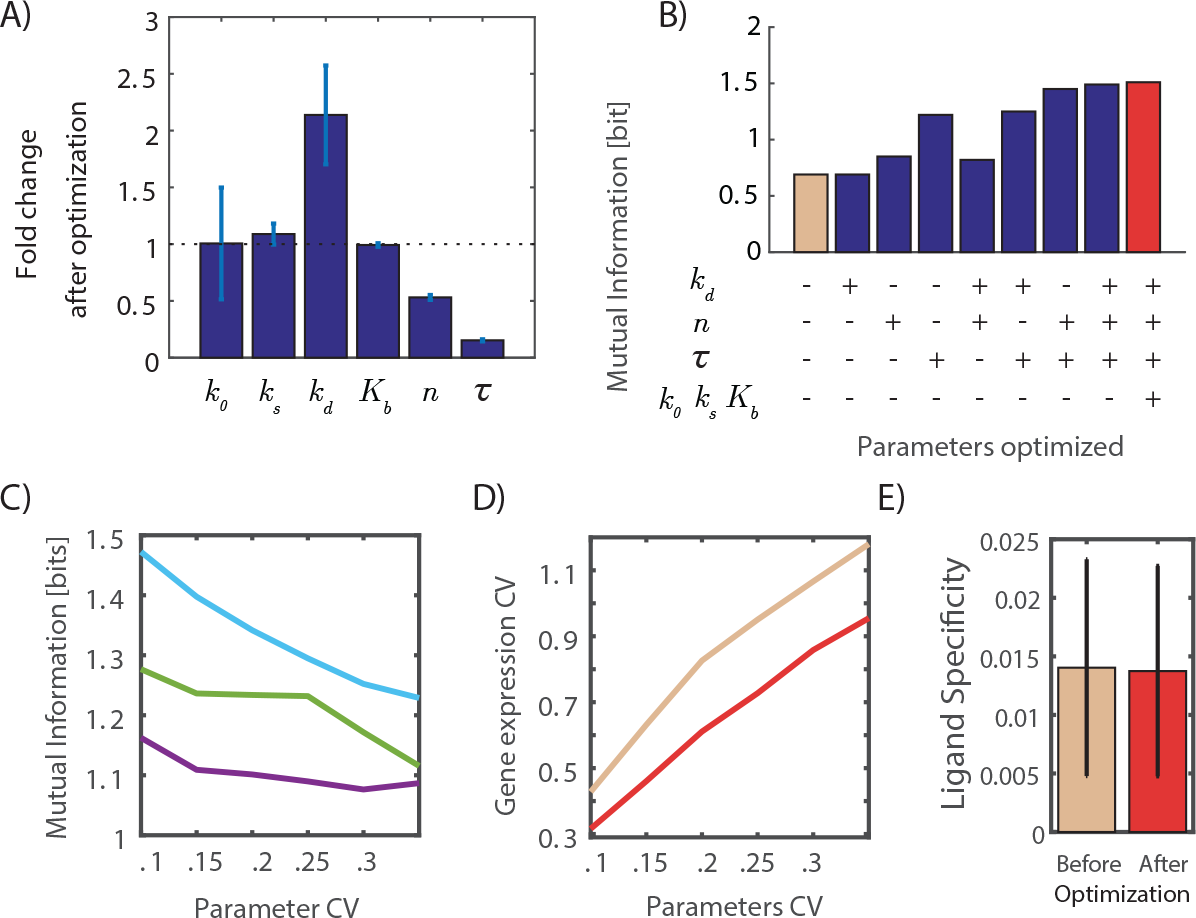
Effects of mutual information optimization on gene expression decoding. **A.** Changes in the six parameters of the decoding model as a result of optimization to increase mutual information. Error bars show standard deviation from 4 repeated optimization runs. **B.** Mutual information with all parameters used at nominal values (light orange), optimized values (red), or a mixture of nominal and optimized values (blue bars). **C.** Mutual information as a function of the amount of variability in a single decoding model parameter shown for mRNA degradation rate k_d_ (purple), hill coefficient n (green) and time delay tau (cyan). **D**. Change in gene expression variability with an increase in encoding and decoding models parameter variability for nominal (light orange) and parameter optimized (red) decoding models. **E** Ligand specificity was calculated as the mean square error between the average responses to the five ligands shown before and after parameter optimization to maximize mutual information.

Next, we examined the contribution of each of the optimal parameters to the increase in decoding accuracy. Our of the system six parameters, only three changed their value in a consistent manner (Figure 4A): mRNA degradation rate (Kd), the cooperativity of the binding reaction (n), and the time lag between increased NfkB in the nucleus to changes in expression (tau). However, the fact that a parameter changed through our optimization does not necessarily guarantee that this change is meaningful and is responsible for the increase in mutual information. To assess the specific contribution of these three parameters we score the decoding accuracy for all possible combination of these parameters for a specific value of variability (25% parameter CV). In each test, all other parameters have their original reference value before optimization. We saw that the parameter that contributed most to the reduced variability is the time delay parameter tau (Figure 4B). This was true for a wide range of degree of biochemical variability. Given the dynamic nature of NfkB, a reduction in the value (and variance) of the time delay unifies the population response and can explain the reduction in gene expression variability we observed (Figure 4D).

Metazoan signaling architecture is unique with a high degree of crosstalk through the network (Rowland *et al*, 2017). The benefits of this architecture are that it is more plastic and allows multicellular organisms to have specialized cells type that responds differently to the same environment without exponentially growing number of pathways (Rowland *et al*, 2017). However, this benefit comes at a cost. Cells need to accurately encode stimulus information in specific signaling patterns that can be decoded by downstream effectors. Biochemical variability in either encoding or decoding will prevent the accurate response to changes in the extracellular environment.

In this study, we analyzed the fidelity of a simple decoding mechanism. We generated realistic signaling patterns using a complex mathematical model and asked how much information about ligand identity is lost in the decoding step. We showed that for realistic patterns of NFkB dynamic gene expression decoding is capable of preserving most of the information that is encoded in the signaling patterns of NFkB.

We focused the decoding question on a single gene model. It is interesting to ask whether more complex patterns of decoding, i.e. multiple genes, can further increase decoding fidelity. To a large degree, this questions depends on the amount of information that exists in the NfkB signaling patterns themselves. Our results show that even a single gene can recover most of that information so while additional genes will show further improvement, it will be bounded by the information loss in the encoding step itself. Where more complex decoding patterns might be important is in allowing decoding parameters that are not optimal. Multiple noisy decoders might increase the overall fidelity over a single noisy one.

Dynamical signaling patterns are observed in multiple signaling systems. Our analysis focused on a single case study of NFkB. It will be important to address questions of decoding fidelity, both computationally and experimentally, in other signaling systems. As our understanding of the role of signaling patterns increases so is the desire to utilize this understanding to design better therapeutics approaches (Behar *et al*, 2013; Bugaj *et al*, 2018). Understanding the mechanism of signal decoding, and the constraints that influence decoding fidelity is an important step in the rational design and manipulation of these dynamical targets.

## Methods

### Quantification of information transduction

To estimate mutual information transduced by NFkB and downstream gene, we applied binless strategy (Selimkhanov et al. 2014). This approach uses an embedding of each simulated temporal response into a vector space and Shannon’s entropy calculates based on the k-nearest neighbor Euclidean distance within these vector spaces. In our calculation, we considered five [50, 100, 150, 300, 450] time points (in minutes) to generate vector space that capturing maximum dynamical feature of NFkB and target gene.

### Stochastic Optimization

Optimization was implemented using simulated annealing (SA) to decipher the optimal decoding parameter set that can transduce maximum information (Optimal Parameter Set = *arg max MI*(*Gene*[*parameters*])). The optimization objective function follows an information-maximization approach between ligand and gene expression. During the simulation, a randomly chosen variable among 6 gene regulation variables was allowed to take a move for each sampling. The maximum step length taken in our simulation is of 5% with respect to the value of the particular variable (in log10 scale) at the prior sampling step. To be explicit, for any parameter (Param), the update was done by the SA rule, Param0= Param + Param × (−1)^n^× δ × rand, where Param0 is the updated value of Param, n is a random integer [0 or 1], δ is the amplitude of allowed change (kept 5%), and rand is a uniform random number between 0 and 1. Using the information of the updated parameter, mutual information is calculated for the new set of vector spaces after each iteration and rejection or acceptance of updated parameter obeys SA rule. With the progress of iteration, the mutual information initially goes up and converges as the output becomes close to the desired value. The mutual information will be optimal if MI value at the i^th^ step of the iteration is close to the i-1^th^ step

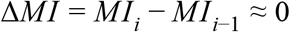

### Parameter Sampling

Model parameters were sampled for the encoding and decoding models (Fig 1AB). For the encoding models, parameters were sampled from a log-normal distribution centered around the reference values with an identical coefficient of variability (CV) for all parameters. For the decoding model, parameters were sampled using a multivariate 6-dimensional uniform distribution for each encoding biochemical variability magnitude. Distribution scales are bounded within biologically relevant parameter range: i.e.,

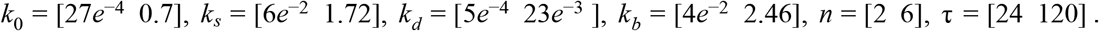

